# Large-scale Visuomotor Reaction Time Self-Testing Reveals Subtle Motor Changes in Older Adults with Subjective Cognitive Impairment

**DOI:** 10.64898/2026.01.02.697425

**Authors:** X. Wang, A.D. Bindoff, R.J. George, E. Roccati, R. Li, K. Lawler, W.M. Connelly, S.N. Tran, A.E King, J.C. Vickers, Q. Bai, J. Alty

## Abstract

**Introduction:** Affordable tools for early Alzheimer’s disease (AD) detection could support drug development and early intervention. Subtle motor changes may indicate preclinical AD, but hand response selection and initiation speeds are understudied. This study assessed whether unsupervised, online visuomotor reaction time (RT) tests relate to subjective cognitive impairment (SCI), a validated high risk state for future conversion to AD

**Methods:** A total of 910 participants (age 66.3±7.5, 70.8% female) completed assessments of simple and choice visuomotor RT tests at home as part of the online TAS Test protocol; they also completed the Cambridge Neuropsychological Test Automated Battery (CANTAB) episodic memory and executive function tests. Among them, 142 participants reported SCI.

**Results:** On the TAS Test visuomotor tests, SCI was associated with 8.4% [1.4%, 15.4%] longer RT ( (*p* = .008; adjusted for task complexity), greater odds of time-out failure (OR = 1.35 [1.01, 1.81]; *p* = .037), and greater variance in RT (log-variance (SCI – comparison) = .094 [.028, .159]; *p* < .001). There were no significant differences between the SCI and comparison groups on any of the CANTAB tests. After adjusting for SCI status, none of the CANTAB tests were significantly associated with RT.

**Discussion:** SCI was associated with longer and more variable visuomotor RT, and greater odds of time-out failure, while not being associated with tests of memory and executive function. Cognitive test scores did not explain a significant amount of variance in visuomotor RT. Taken together, these results support a hypothesis that people with SCI may be experiencing earlier visuomotor deficits that are distinct from (or precede) decline in episodic memory and executive function. Visuomotor tasks that record RT may be more sensitive to preclinical manifestations of cognitive decline than more traditional tests of cognitive function.

## 1 Introduction

It is estimated that dementia affects 55 million people globally, with numbers set to triple by 2050 [1]. Alzheimer’s disease (AD) is the most common cause [2], marked by a 10-20 year presymptomatic phase of pathology [3]. Within the older population, a stage of subjective cognitive impairment (SCI) often precedes a dementia diagnosis by many years. During this stage, individuals may feel that their cognitive abilities are not what they used to be, yet they still perform well on standard cognitive tests. As a result, a ‘wait and see’ approach is typically adopted. Identifying individuals at-risk for AD, such as those with SCI [4–8] , is key to early intervention and risk factor modification which may delay or prevent up to 45% of cases [9]. However current cognitive tests often lack the sensitivity to detect subtle, early changes, and performance may be masked in individuals with higher levels of education [10]. This underscores the need for more sensitive measures that capture early brain alterations associated with AD.

New monoclonal therapies for AD are anticipated to be most effective when administered earlier in the disease process [11, 12]. However, the absence of affordable and accessible tests presents a significant challenge to dementia prevention, as current methods to detect the earliest stages of AD pathology - such as neuropsychological assessments, blood and cerebrospinal fluid (CSF) biomarkers, and positron emission tomography (PET) scans - are often too expensive, time-consuming, or invasive for widespread use [13]. Compounding this issue is a global shortage of diagnostic imaging capacity [14]. The rising demand for early, and at times speculative, neuroimaging, risks overwhelming already strained healthcare systems, particularly since monoclonal antibody therapies require ongoing monitoring through MRI [15]. Given that the majority of individuals with dementia are expected to reside in low- and middle-income countries (LMICs) [16], there is an urgent need for cost-effective, home-based testing solutions to enable early detection.

Hand motor function is emerging as a promising biomarker due to its low cost and accessibility through widely available computer technologies, such as video recording on laptops and smartphones [17]. Drawing from gait literature, we know that motor patterns have demonstrated significant potential in identifying individuals at risk of AD [18, 19]. Gait anomalies, characterised by slower and less rhythmic stepping, have been found to emerge up to a decade before diagnosis of dementia [18]. Findings from our own research on repetitive hand movements, such as keyboard tapping, have indicated that motor characteristics such as slower tapping speed, greater variance, longer dwell time on the keys, and reduced accuracy, are associated with lower episodic memory performance in asymptomatic individuals [20]. This not only provides construct validity, but it is also particularly relevant to AD as episodic memory decline is typically one of the earliest cognitive changes [21]. Findings from our own research on repetitive hand movements, such as webcam-based finger tapping, indicate that motor characteristics extracted from these brief tests improve prediction of episodic memory, executive function, and working memory in cognitively asymptomatic older adults. These results suggest that subtle variations in tapping behaviour may serve as low-cost, remote markers for identifying individuals at elevated risk for cognitive decline [17]. Gait and key-tapping metrics were moderately correlated, but each showed distinct associations with cognitive impairment. Combining these motor measures substantially improved classification of dementia and MCI beyond gait alone or demographic factors, indicating that gait and key-tapping capture complementary aspects of motor–cognitive dysfunction [22]. Hand motor dysfunction was evident across the dementia continuum, with key-tapping frequency, speed, and rhythm strongly associated with memory and executive performance and with diagnoses of SCI, MCI, and dementia. Key-tapping measures accurately differentiated dementia and MCI from healthy controls, suggesting that hand movement analysis may offer a promising, accessible motor biomarker for cognitive impairment pending further validation [23]. Smartphone-based analysis of hand and speech-like movements, such as the TapTalk test, may detect subtle motor changes associated with preclinical Alzheimer’s disease long before cognitive symptoms emerge. A large multi-phase validation program is underway to determine which motor features best predict AD biomarkers and clinical diagnoses, with the goal of enabling low-cost, population-level screening for early AD risk [24]. Upper-limb motor assessments—including finger tapping, pegboard tasks, and functional actions like writing—are frequently used in studies of cognitive impairment, though protocols vary widely. Across 60 studies, slower speed, more errors, and greater movement variability were generally associated with cognitive impairment, highlighting both the potential of upper-limb motor function as a biomarker and the need for more standardised assessment methods [25].

Building upon basic hand motor tasks involving repetitive hand movements, such as keyboard tapping [20, 23] and finger-to-thumb tapping [17, 24], a logical next step for deepening the investigation is to incorporate more complex motor-cognitive assessments, such as visuomotor reaction time tests (RTT). In these tasks, a movement executed in response to a visual stimulus with no need for response selection, apart from the timing of the response, is referred to as simple reaction time (SRT). In contrast, when the task requires selecting between multiple possible responses - in which stimulus features are used to determine the direction of movement or which limb (e.g., left or right hand) to use - it is known as choice reaction time (CRT). SRT and CRT tasks allow for a broader exploration of sensorimotor integration and cognitive processing, potentially offering greater sensitivity to subtle impairments. Several small studies found that people with AD, or probable AD, have prolonged SRT and greater variance of SRT and CRT compared to healthy controls [26–34]. For example, Müller et al. found that in 21 participants with mild AD, there was a significantly prolonged SRT (149%) compared to 15 healthy age-matched controls when 10 repetitions were completed using a response key [29]. More recently, a 2018 study involving 23 individuals with possible/probable AD and 25 healthy controls, identified greater variance in RT among the AD group for both SRT and CRT, when using the computerised Cambridge Neuropsychological Test Automated Battery (CANTAB) [27]. Both studies were limited by small sample sizes and were conducted in controlled research settings, but they suggest the potential utility of RTTs as more sensitive markers of cognitive-motor dysfunction in early AD.

Several knowledge gaps remain regarding the association between RTTs and cognitive changes, primarily due to the limitations of previous studies [29], which often lacked adjustments for potential confounders, and did not focus on earlier stages of the AD continuum. Particularly underexplored are “at-risk” groups, such as individuals with SCI. Furthermore, most studies have been conducted in small clinical cohorts under researcher supervision, utilising specially designed microswitches or response boxes to measure RT. Notably, no studies to date have examined the association between remotely administered RTTs and cognitive function assessed through online platforms. This limitation restricts both scalability and translational potential. The ability to administer RT tasks online opens the possibility of broader access, particularly for individuals in rural or underserved areas with limited clinical resources.

TAS Test is a self-administered online motor-cognitive battery developed to detect early impairments seen in AD. TAS Test has been used remotely and been found to be feasible and acceptable in > 2300 older adults regardless of age, education and computer literacy [35, 36]. TAS Test includes a visuomotor RTT protocol, with measures of SRT, CRT and action time (AT). SRT and CRT measure the interval between the appearance of a visual stimulus and the initiation of the action, whereas AT captures the period between the initiation of the action to the completion of the movement to the target. We hypothesised that slower visuomotor RT and AT would be associated with more errors on cognitive tests in asymptomatic older adults, and with SCI, which is not typically associated with poor cognitive test scores (adjusted for age and sex). We further hypothesised that greater variance in visuomotor RT and AT, and a higher probability of time-out failure, would also be associated with lower cognitive test scores and SCI.

## 2 METHODS

### 2.1 Ethics and consent

The Human Research Ethics Committee of the University of Tasmania approved the TAS Test Project (ref. HREC H0021660; registered on the ClinicalTrials.gov registry as NCT05194787) and the ISLAND Project (ref. HREC H40267). All participants gave informed consent.

### 2.2 Study participants

Participants were selected from a community-based cohort of individuals aged 50 years and older living in Tasmania, Australia. These individuals were enrolled in ISLAND (Island Study Linking Ageing and Neurodegenerative Disease, a public health initiative with a primary focus on educating participants about modifying dementia risk factors [37]. Described in detail by Bartlett et al., the ISLAND protocol allows the annual collection of participant data through self-reported surveys comprising demographics as well as information on diagnoses such as Parkinson’s disease (PD), Multiple Sclerosis (MS) and dementia [37]. Between June and August 2021, participants in ISLAND were invited via email to take part in TAS Test which involved a comprehensive online battery of motor-cognitive assessments, including simple and choice RTT, administered through a dedicated website (see Alty et al. for a detailed protocol) [36].

Individuals who responded “yes” to either of the following questions in the ISLAND annual survey were excluded from the analysis, as our study does not focus on individuals with objective subjective impairments: 1. “Have you been told by a doctor that you have dementia?” 2. “Have you been told by your doctor that you have memory impairment but you are uncertain if you have dementia?”. Of the remaining individuals, two groups were examined: the SCI group, consisting of those who answered “yes” to the following question, and the asymptomatic group, comprising those who answered “no”: “Have you noticed a substantial change in your memory and mental function in recent years?” Additionally, individuals with diagnoses of PD and MS were excluded from the cohort. This single question approach to define SCI is a validated method, supported by longitudinal evidence [7, 38].

### 2.3 Reaction time tests

RTTs were conducted as part of the TAS Test protocol [36]. Designed for self-administration at home, TAS Test collects time-stamped data for each test. The test is self-paced, featuring an instruction page, and a practice session to ensure that participants understand the protocol. This protocol includes a simple visuomotor RTT followed by a choice RTT. Participants complete the test using a laptop or desktop computer, employing a computer mouse or a trackpad. The protocol included two practice trials followed by five test trials for both the simple and choice RTT. Participants were asked to indicate their hand dominance (“What hand do you usually write with?”) and were guided through the tasks via on-screen instructions.

In the simple RTT, a single white circle is displayed toward the top of the screen, and participants are instructed to begin a trial by moving the mouse cursor to the “HOLD” button located at the bottom of the screen, and press and hold the left mouse button (see Figure 1A). This action triggers the circle to change from white to yellow after a randomised delay of 2, 3, or 4 seconds. Participants are then instructed to release the mouse button, move the cursor to the yellow circle, and click on it as quickly as possible. In the choice RTT, the initial “HOLD” button action and the random delay waiting for the target stimulus is the same as the SRT tasks, however in this test, one of five white circles displayed at the top of the screen randomly turns yellow, and participants are instructed to click on the circle that changes to from white to yellow as quickly as possible (Figure 1B).

**Figure 1.**
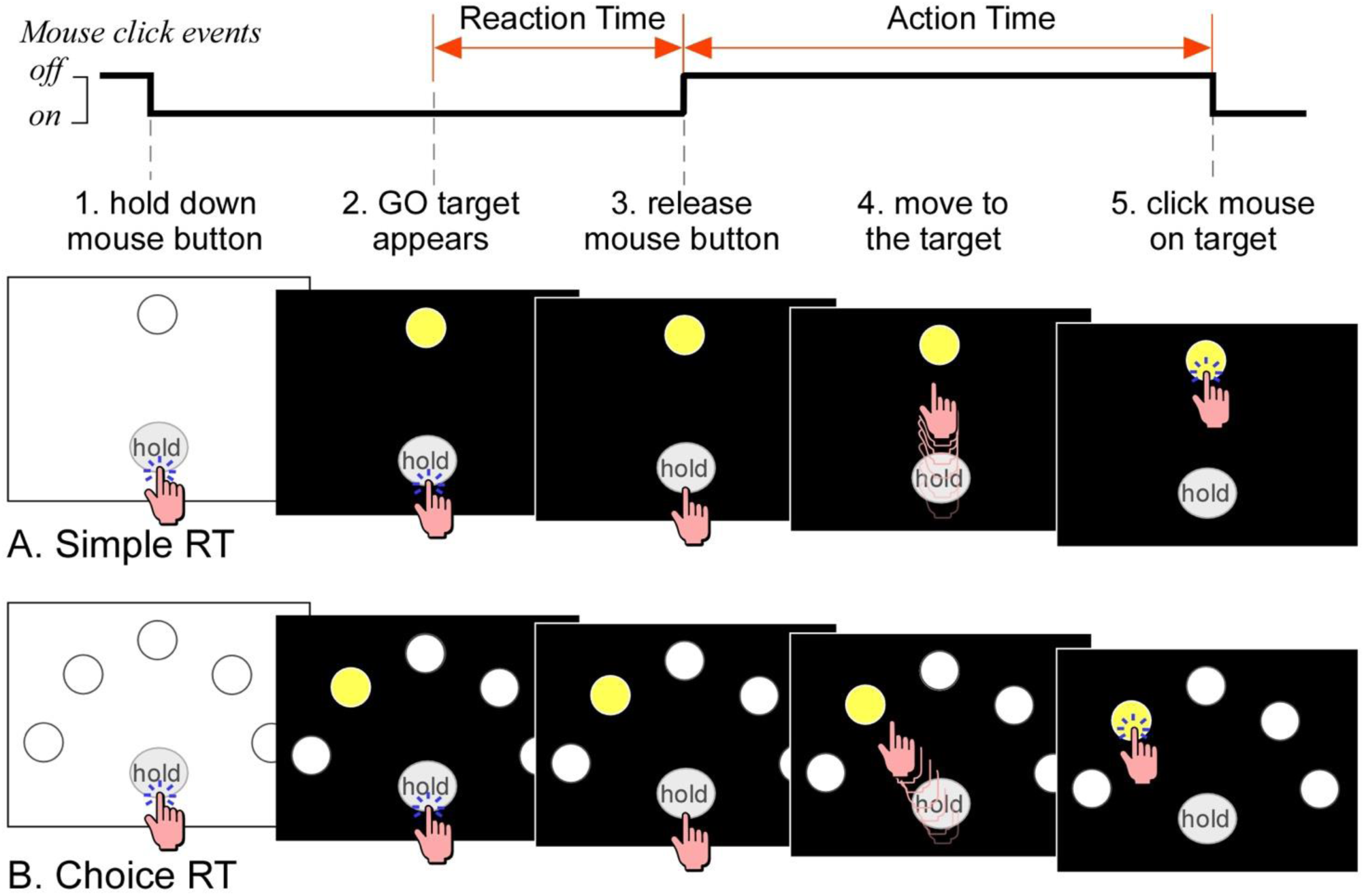
Schematic diagram depicting the calculation of reaction time (RT) and action time (AT) in both simple (A) and choice reaction time (B) tests. RT is defined as the interval between the appearance of the GO target (yellow circle) and the release of the mouse button on the “HOLD” button. AT is the latency from the release of the mouse button to the moment the target is clicked.

In the simple RTT, there is spatial certainty about the movement trajectory and end point, and the task predominantly engages attention and sensorimotor functions. In contrast, for the choice RTT, the response cannot be preplanned and requires additional cognitive processes, including decision-making and motor planning. In August 2021, participants also completed the Paired Associates Learning (PAL) and Spatial Working Memory (SWM) tasks from CANTAB, a validated cognitive function assessment battery that provides objective measurements [39].

The timing of the visual stimuli is synchronised with the screen refresh, ensuring minimal delay to display, thereby allowing for accurate measurement of RT via recorded mouse clicking events. In order to confirm the accuracy of reported reaction times, an ATmega32u4 microcontroller sampled the white circle via a photodiode, and when it changed to yellow, after a random delay (0 to 300 ms), a click was sent via USB to the computer. The microcontroller recorded the duration from the stimulus change to when it sent the click. This was repeated 20 times. These times as reported to by TAS Test and the microcontroller were compared via linear regression. The R^2^ of this relationship was 0.98 with a slope of 0.997 ± 0.04, meaning that the relative changes in reaction time are extremely accurate. The intercept of this relationship, however, was 118 ± 6 ms, meaning that TASTEST reported times are approximately 120 ms slower than the microcontroller time. The source of this error is likely due screen latency, that is, the TASTEST timer starts when the stimulus is requested to be drawn, rather than when it appears on screen. However, this was not investigated further. All times reported in the manuscript are those reported by TASTEST and were not adjusted for potential display errors.

The same procedures for calculating RT and AT are applied in both tests, as illustrated in Figure 1.

### 2.4 Cognitive performance

Participants completed a subset of the CANTAB battery, which is an online cognitive test assessing episodic memory, working memory and executive function. CANTAB is a validated tool with strong test-retest reliability [40] and is commonly used to assess age-related cognitive decline, including in conditions such as AD [41].

#### 2.4.1 Episodic memory assessment

The PAL test from CANTAB evaluates episodic memory and typically takes about 8 minutes. As shown in Figure 2, the boxes are “opened” in a randomised order, with one or more boxes containing a pattern. Participants are required to remember the pattern displayed behind the boxes (Figure 2A and 2B). PALTEA6 represents the total errors made at the six-pattern stage, adjusted for incomplete or failed tests.

**Figure 2:**
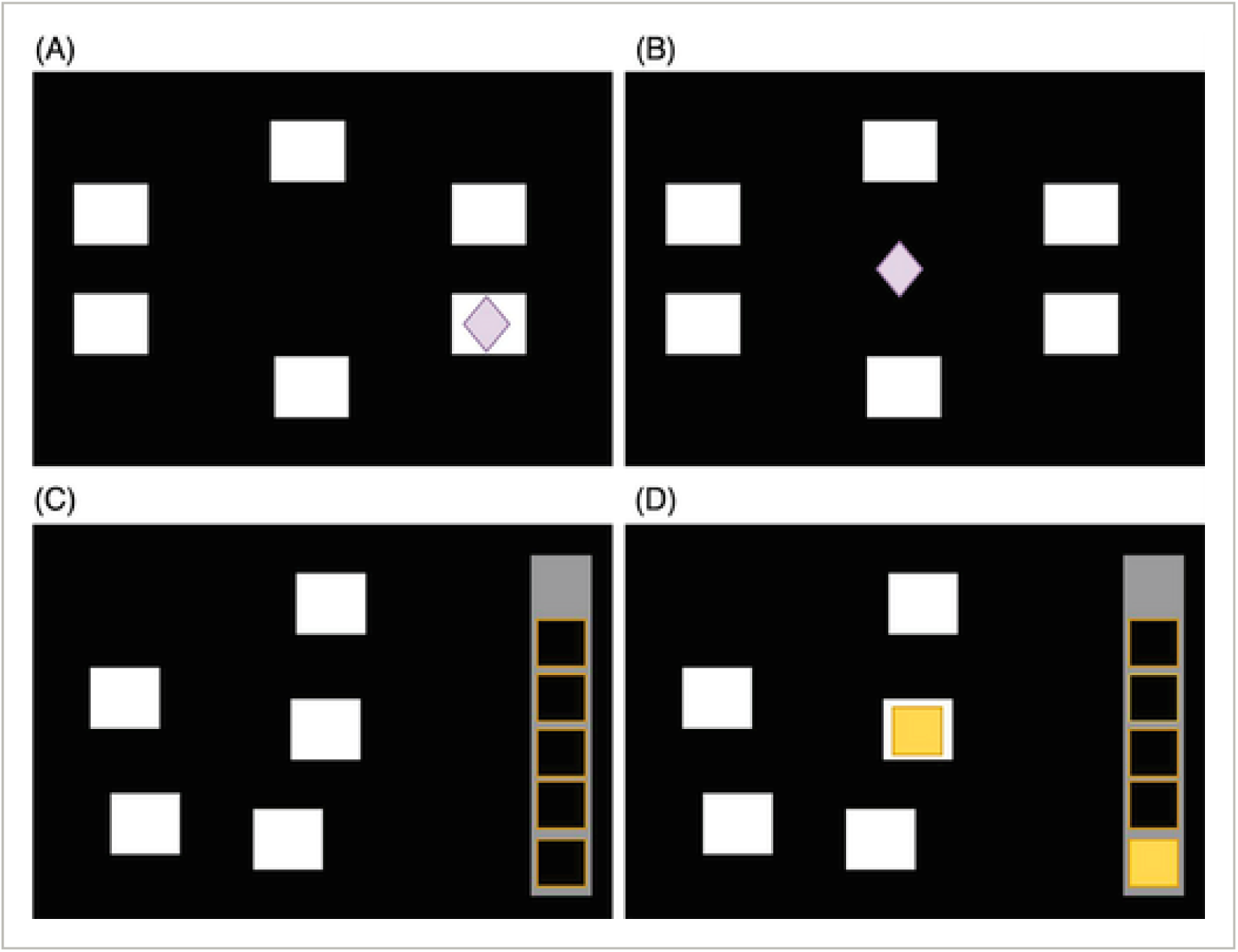
A schematic diagram of the stages of the Paired Associates Learning (PAL) and Spatial Working Memory (SWM) cognitive tests. For PAL, (A) participants were required to remember different patterns “stored” in various boxes on the screen. (B) A new pattern would appear at the centre of the screen, and participants were asked to select the box where the same pattern was stored. For SWM, (C) and (D), participants were required to click on the boxes to identify where yellow tokens were stored. Yellow tokens could only appear once in one box per attempt. In each attempt, there could be three, four, or six boxes which had yellow tokens.

#### 2.4.2 Working memory and executive function assessment

The SWM test evaluates aspects of executive function, including the working memory component. In this task, participants try to locate a single yellow “token” among several boxes using a process of elimination (Figure 2C and 2D). In a single attempt, only one “token” is hidden in a single box. The number of boxes gradually increases to a maximum of 12, and in each attempt, participants click on the boxes to find the “token” until the empty column on the right-hand side of the screen is filled with “tokens”. Performance can be measured by two key outcomes: *Spatial Working Memory Between Errors at 6 boxes* (SWMBE6), which reflects the number of times a participant re-clicks a box that has already been selected (an index of working memory), and *Spatial Working Memory Strategy* (SWMS), which captures the number of different boxes used to begin each new search (an index of strategic planning, an executive function process).

### 2.5 Data analysis

RT of mouse clicks were recorded in milliseconds (ms), with time-out failure defined as an RT exceeding 2000ms for a given trial (i.e., failing to release the “HOLD” button within 2 seconds after a circle turns yellow). The two practice trials from the simple and choice RTT were excluded from the analysis, leaving five trials of each per participant. Simple and choice RTT were identified by a dummy variable, with the data arranged in long-format for mixed effects analysis.

Errors on CANTAB tests were recorded as counts, and a dummy variable was created to denote participants with self-reported subjective cognitive decline (no formal diagnosis). Age (years) was mean-centred and standardised for computational benefit. Sex was recorded as “male” or “female” according to participant response. RTs greater than 2000ms were recorded as zeros in the raw data to denote time-out failure on that trial (more on how these were handled below). Reaction and action times <150ms were excluded as implausible.

To relax the assumption of independence we used mixed effects regression models, fitting by-participant random intercepts and random coefficients for the choice task (allowing participant-level random effects to vary between simple and choice tasks). Specifically, we used the ‘glmmTMB’ package [42] in R [43] to estimate Gaussian hurdle models which assume zeros (time-out failures in our case) are produced by a different process to the valid times and occur with some binomial probability, which can be estimated via the logit-link function for fixed effects parameters. We applied log-transformation to RT+1 such that time-out failures (recorded as zero in the raw data) remained zero and the non-zero RTs were approximately normally distributed. We log-transformed AT+1 similarly.

The ‘glmmTMB’ package further allows the dispersion parameter (in the Gaussian hurdle case, log-variance) to be conditioned on fixed effects, which not only allows the assumption of homogeneity of variance to be relaxed, but also tests the null-hypothesis that dispersion does not vary with whichever fixed effects are included.

Making use of the flexibility offered by ‘glmmTMB’ in estimating the joint distribution of a conditional model, we fitted a fixed-effects interaction between SCI and task complexity (SRT, CRT) regressed on RT and AT (in separate univariate models), adjusted for age and sex. The dispersion component estimated log-variance for comparison and SCI groups. For the RT models, the hurdle component estimated the log-odds of time-out failure conditioned on SCI. To estimate the effects of each CANTAB variable, we then used these same models but replaced the interaction, dispersion, and hurdle terms with CANTAB variables, and adjusted for the fixed effect of SCI. R code is provided in the Supplementary Materials. We report (adjusted) Wald estimates for p-values and 95% confidence intervals (CI). Fixed effect planned contrasts for interaction terms report t-statistics. All 95% CI and significant p-values are adjusted using the Šidák correction for a family of 8 comparisons (2 dependent x 4 independent variables) to maintain a family-wise Type 1 error rate < .05.

To estimate the association between SCI and cognitive test performance adjusted for age and sex, we used Poisson regression to regress SCI, age and sex on CANTAB variables. We report the results of likelihood ratio tests.

To assess fit of Gaussian hurdle models we used posterior-predictive plots. For other models we used the appropriate graphical assumption checks (residual plots, Q-Q plots for normality of residuals and random effects, *etc*).

## 3. RESULTS

A total of 945 participants who were eligible from the ISLAND cohort (n = 2766) completed CANTAB cognitive tests and TAS Test RTT. Table 1 outlines the demographic data of the cohort. The number of people excluded from the analysis due to affirming or not responding to at least one of the following questions: 1. ‘Have you been told by a doctor that you have dementia?’ (n = 16). 2. ‘Have you been told by your doctor that you have memory impairment but are uncertain if you have dementia?’ (n =9). Participants who indicated they have PD (n = 6) and MS (n = 4) were excluded from the study, leaving a total of 910 participants in the analysis. Of these, 142 answered “yes” to the SCI screening question (Table 1).

**Table 1:** Demographic Data, Reaction Time and Action Time of the Cohort.

|  | No reported change<br>in recent memory or<br>mental function<br>(N=768) | SCI*<br>(N=142) |
| --- | --- | --- |
| **Sex** |  |  |
| Female | 548 (71.4%) | 96 (67.6%) |
| Male | 220 (28.6%) | 46 (32.4%) |
| **Age (years)** |  |  |
| Mean (SD) | 66.1 (7.55) | 67.1 (7.45) |
| Median [Min, Max] | 66.0 [51.0, 90.0] | 67.5 [53.0, 84.0] |
| **Trial time-out failures** |  |  |
| Mean (SD) | 0.556 (0.854) | 0.704 (1.06) |
| Median [Min, Max] | 0 [0, 4.00] | 0 [0, 5.00] |
| **Reaction time simple (ms)** |  |  |
| Mean (SD) | 639 (233) | 698 (256) |
| Median [Min, Max] | 558 [348, 1760] | 617 [311, 1640] |
| **Reaction time choice (ms)** |  |  |
| Mean (SD) | 679 (254) | 777 (325) |
| Median [Min, Max] | 585 [382, 1800] | 638 [389, 1920] |
| **Action time simple (ms)** |  |  |
| Mean (SD) | 661 (179) | 655 (188) |
| Median [Min, Max] | 654 [234, 1710] | 620 [212, 1120] |
| **Action time choice (ms)** |  |  |
| Mean (SD) | 678 (184) | 674 (228) |
| Median [Min, Max] | 685 [211, 1240] | 697 [224, 1240] |
| **Episodic memory (PALTEA6)** |  |  |
| Mean (SD) | 4.56 (4.79) | 4.85 (5.55) |
| Median [Min, Max] | 3.00 [0, 22.0] | 3.00 [0, 20.0] |
| **Working memory (SWMBE6)** |  |  |
| Mean (SD) | 3.28 (3.30) | 3.49 (3.65) |
| Median [Min, Max] | 3.00 [0, 13.0] | 2.00 [0, 12.0] |
| **Executive function (SWMS)** |  |  |
| Mean (SD) | 7.74 (2.81) | 7.83 (2.70) |
| Median [Min, Max] | 8.00 [2.00, 13.0] | 8.00 [2.00, 13.0] |

\* SCI group answered "yes" to the question "Have you noticed a substantial change in your memory and mental function in recent years?"

### 3.1 CANTAB cognitive test scores and SCI

As expected, there were no significant differences between SCI and comparison groups on any of the CANTAB tests of episodic memory (*p* = .578), working memory (*p* = .895), or executive function (*p* = .975; all adjusted for age and sex).

### 3.2 Visuomotor reaction time and SCI

Participants in the SCI group had significantly longer mean RTs than the comparison group, *p* = .008. Adjusting for the main effect of complexity (but for simplicity, not the interaction between SCI and complexity) this difference was an increase of 8.4% [1.4%, 15.4%] over comparison group RTs. RT was significantly greater in the CRTT (*p* < .001), however the interaction between complexity and SCI was not statistically significant (*p* = .139). Figure 3 shows the estimated means from the conditional model averaged over age and sex.

**Figure 3.**
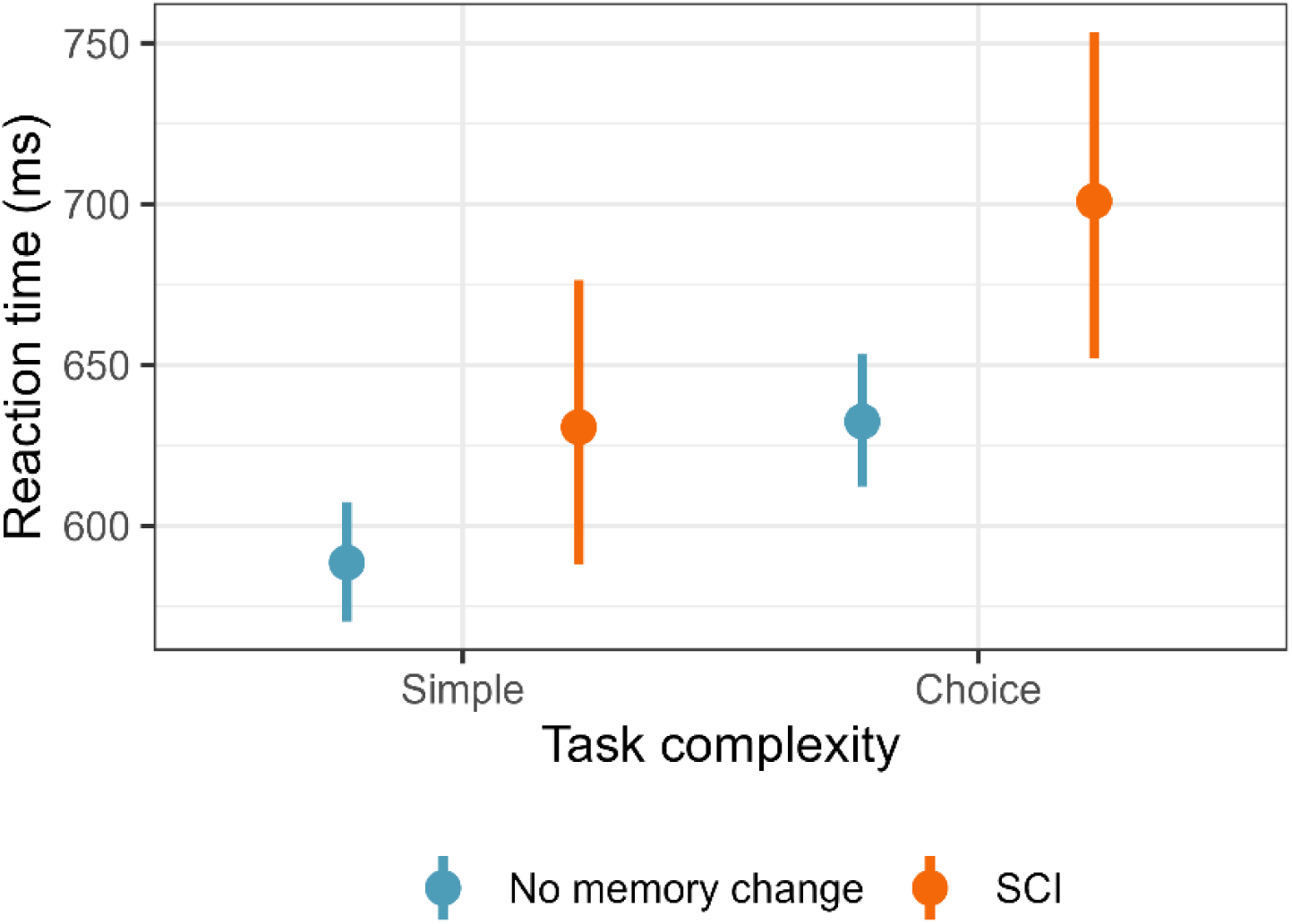
Estimated mean reaction time and Šidák corrected 95%CI for SCI and comparison groups in the SRTT and CRTT, adjusted for age and sex, probability of time-out failure and heterogeneity of variance between groups.

Time-out failures were significantly more likely for participants in the SCI group, OR = 1.35 [1.01, 1.81]; *p* = .037. Log-variance was also significantly greater in the SCI group (mean difference = .094 [.028, .159]; *p =* <.001).

### 3.2 Visuomotor action time and SCI

Log-variance was significantly greater in the SCI group (mean difference = .078 [.029, .127]; *p* = .014), however the difference in AT between groups was not significant, *p* = .262 for the main effect of SCI and *p* = .745 for the interaction between SCI and task complexity.

### 3.3 Visuomotor reaction/action time and cognitive test scores

No CANTAB cognitive tests were associated with visuomotor RT or AT, however each additional error on the executive function task (SWMS) was associated with a .024 [.003, .046] increase in log-variance in RT (*p* = .039).

## 4 DISCUSSION

This study found an association between SCI and prolonged time to make an initial response to a visual stimulus (compared to controls), along with higher odds of time-out failure and greater variance in RTs. The hypothesis that more errors on episodic memory, working memory and executive function tests would be associated with prolonged RTs, time-out failure, and variance in RTs (adjusting for SCI) was not supported, nor was SCI predictive of errors on these cognitive tests. Taken together, these results suggest a link between self-reported changes in memory and cognitive function and visuomotor RT that is not predicted by performance on cognitive tests.

It is also noteworthy that all tests were self-administered and to the best of our knowledge this is the largest RTT study in older adults in the literature. In summary, our findings supported the first hypothesis, that self-reported poorer cognitive function (i.e., individuals with SCI) would be associated with longer RT. However, the second hypothesis, regarding an association between variance in RTs and cognitive function, was not supported. Additionally, poorer cognitive function on cognitive tests (specifically, CANTAB working memory performance) was associated with test failure.

Prior clinical research has examined the relationship between RT performance and the early stages of AD. Müller et al. observed that individuals with mild AD exhibited significantly prolonged RTs, which defined as the time to detect a stimulus and click on the response key, compared to healthy controls in a small clinic-based cohort [29]. Building upon this smaller cohort study, our study, which included over 900 participants, identified associations between long RTs and SCI, which is a risk factor for later AD, and greater dispersion of RTs with poorer cognitive performance, including lower scores in episodic memory, working memory, and executive function. Moreover, Müller et al. employed specialised laboratory equipment for their RT measurements, which may not be universally accessible. In contrast, our study administered the RTT via an online platform, accessible to individuals with laptops or desktops, thereby eliminating the need for in-person attendance. A more recent study from 2018 also found greater variance in RT among 38 individuals with probable or possible AD (n= 23) compared to controls (n= 15) [27]. In our cohort, we examined variance in a large community cohort, finding that variance did not show significant correlations on poorer cognitive function with RT or AT. To our knowledge, this study is the first to investigate both AT and RT in the same context, exploring their associations with cognitive performance. By examining RT (stimulus detection and first reponse) and AT (motor response execution) separately, we can more precisely understand how these components may be independently affected by underlying neuropathology.

Our study offers several notable strengths. First, it represents the first administration of visuomotor RT tests to a large community cohort, including individuals with SCI and an asymptomatic group. The online format of the study facilitates remote participation, overcoming geographic limitations and has the potential to reach a wider population. Second, the design of TAS Test allows for the simultaneous calculation of both RT and AT within the same test, reducing the need for additional assessments and minimising the burden on participants. Additionally, we employed CANTAB, a validated online cognitive assessment tool that evaluates multiple cognitive domains, including episodic memory, working memory, and executive function.

However, this study also has certain limitations. The cohort lacks ethnic diversity, as it is predominantly composed of White participants with Northern European ancestry from Tasmania, Australia [44]. Furthermore, more than 70% of participants are female, and the sample is highly educated, reflecting a common trend in research studies. Additionally, handedness was assessed through a single question regarding left-handed, right-handed, or ambidextrous status, which may not fully capture the complexity of hand dominance. Furthermore, the nature of SCI remains uncertain, as it may indicate early-stage AD, mood disorders such as depression, or other conditions. Finally, a number of potentially confounding variables were not measured in the study, including mood, sleep quality, physical discomfort (e.g., hand pain), visual acuity, caffeine intake, and computer literacy, all of which may influence RT and AT performance.

In conclusion, this study involved a large community cohort, including individuals with SCI, and utilised validated cognitive tests to assess various cognitive domains in the asymptomatic group. By extracting both RT and AT measures from the tests, we identified significant associations between these measures and SCI, which is associated with an increased risk of dementia [45]; our findings suggest that this simple, accessible online motor test may have utility in early risk detection for dementia even when cognitive tests remain objectively normal. Future studies could incorporate a more nuanced approach to adjusting for handedness, using the abbreviated Edinburgh Handedness Questionnaire as this has been integrated into TAS Test [46]. Identifying individuals at high risk of dementia through non-invasive, scalable methods could facilitate earlier intervention and improve recruitment for clinical trials targeting preclinical stages of cognitive decline.

## Acknowledgements

The authors are grateful for the support of the ISLAND project team.

## Data availability statement

The authors may share de-identified data with qualified investigators whose proposal of data use has been approved by an independent review committee.

## Author contributions

Drafting or revision of manuscript: XW, RSG, ADB, RL, WMC, ER, QB, AK, JV, JA, ST, KL.

Study concept or design: JA, RSG, ER

Acquisition of data: XW, RL, ER, WMC, JV, AK, QB, JA, KL.

Analysis or interpretation of data: XW, ADB, RSG, RL, ER, AK, JV, JA.

## Supplementary Materials

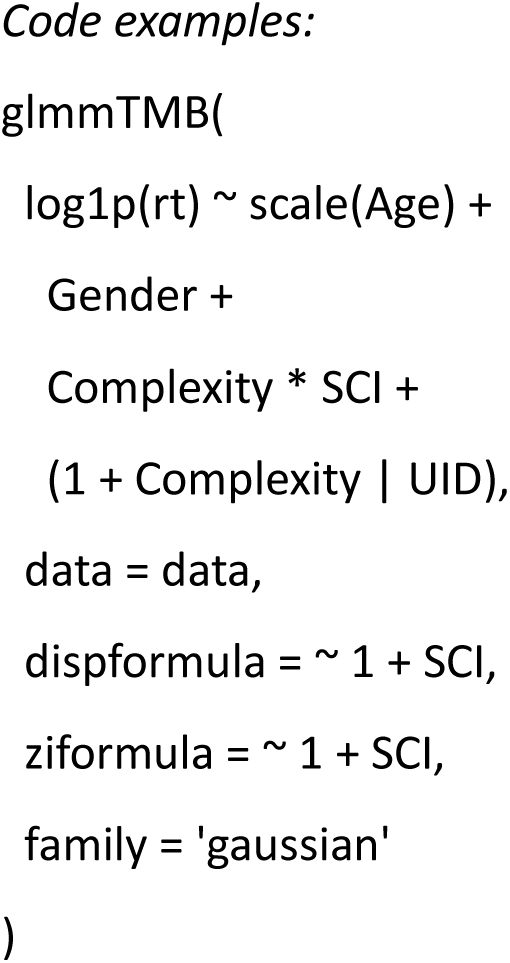
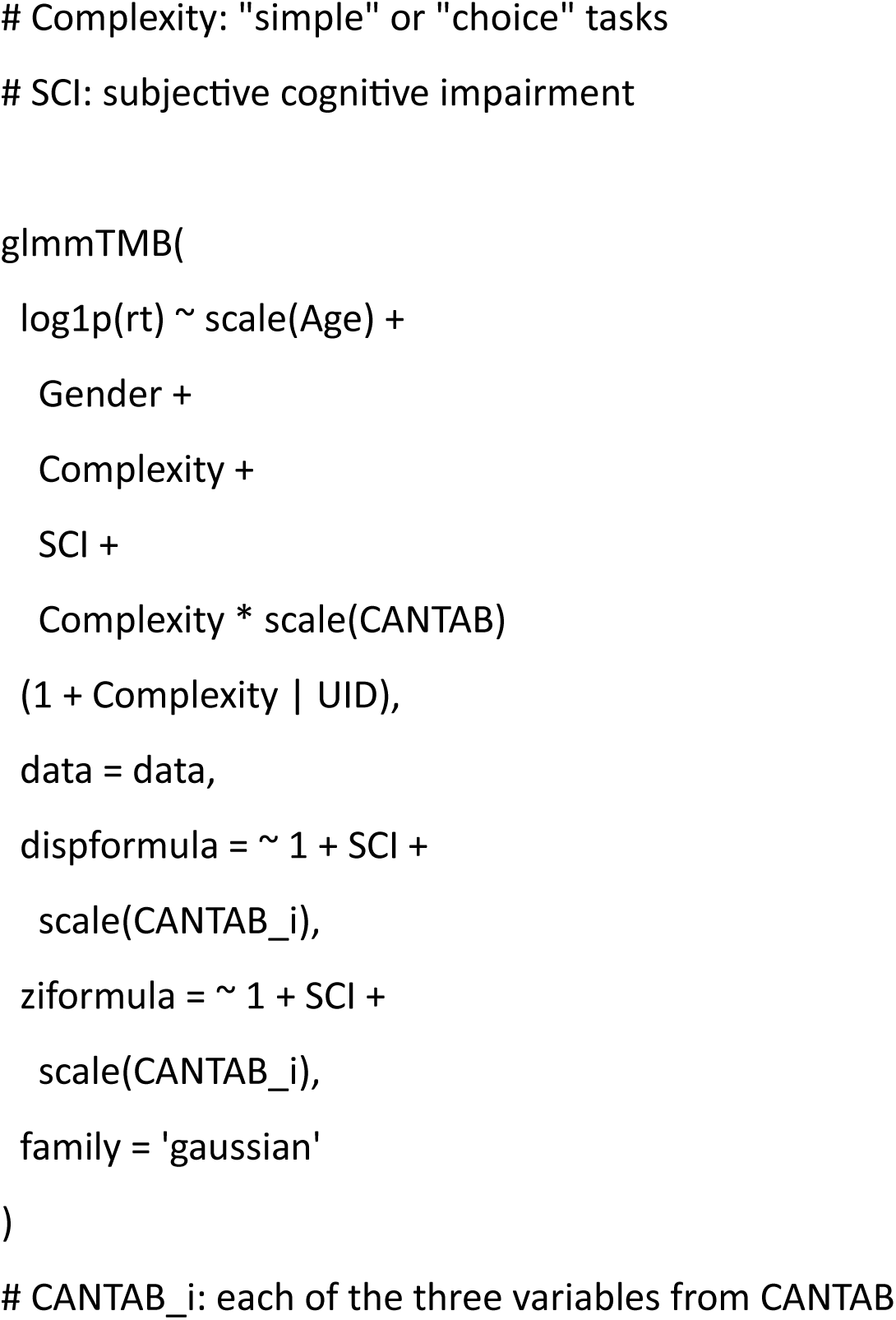

## References

1. Livingston, G., et al., The Lancet International Commission on Dementia Prevention and Care. Lancet, 2017. 390(10113): p. 2673–2734.

2. World Health Organization, Global action plan on the public health response to dementia 2017–2025. 2017.

3. Sperling, R.A., et al., Toward defining the preclinical stages of Alzheimer’s disease: Recommendations from the National Institute on Aging-Alzheimer’s Association workgroups on diagnostic guidelines for Alzheimer’s disease. Alzheimer’s & dementia, 2011. 7(3): p. 280–292.

4. Garcia-Ptacek, S., et al., Subjective cognitive impairment: Towards early identification of Alzheimer disease. Neurología (English Edition), 2016. 31(8): p. 562–571.

5. Montejo, P., et al., Subjective memory complaints in the elderly: prevalence and influence of temporal orientation, depression and quality of life in a population-based study in the city of Madrid. Aging & mental health, 2011. 15(1): p. 85–96.

6. Semba, R.D., et al., Motoric cognitive risk syndrome: Integration of two early harbingers of dementia in older adults. Ageing research reviews, 2020. 58: p. 101022.

7. Pike, K.E., et al., Subjective cognitive decline: level of risk for future dementia and mild cognitive impairment, a meta-analysis of longitudinal studies. Neuropsychology review, 2022. 32(4): p. 703–735.

8. Mitchell, A.J., et al., Risk of dementia and mild cognitive impairment in older people with subjective memory complaints: meta-analysis. Acta Psychiatrica Scandinavica, 2014. 130(6): p. 439–451.

9. Livingston, G., et al., Dementia prevention, intervention, and care: 2024 report of the Lancet standing Commission. The Lancet, 2024. 404(10452): p. 572–628.

10. Meng, X. and C. D’arcy, Education and dementia in the context of the cognitive reserve hypothesis: a systematic review with meta-analyses and qualitative analyses. PloS one, 2012. 7(6): p. e38268.

11. Livingston, G., et al., Dementia prevention, intervention, and care: 2020 report of the Lancet Commission, in The Lancet. 2020, Lancet Publishing Group. p. 413–446.

12. Van Dyck, C.H., Anti-amyloid-β monoclonal antibodies for Alzheimer’s disease: pitfalls and promise. Biological psychiatry, 2018. 83(4): p. 311–319.

13. Li, R., et al., Applications of artificial intelligence to aid early detection of dementia: a scoping review on current capabilities and future directions. Journal of biomedical informatics, 2022. 127: p. 104030.

14. Jeganathan, S., The growing problem of radiologist shortages: Australia and New Zealand’s perspective. Korean Journal of Radiology, 2023. 24(11): p. 1043.

15. Lacorte, E., et al., Safety and efficacy of monoclonal antibodies for Alzheimer’s disease: a systematic review and meta-analysis of published and unpublished clinical trials. Journal of Alzheimer’s Disease, 2022. 87(1): p. 101–129.

16. Mukadam, N., et al., Changes in prevalence and incidence of dementia and risk factors for dementia: an analysis from cohort studies. The Lancet Public Health, 2024. 9(7): p. e443–e460.

17. Li, R., et al., Brief webcam test of hand movements predicts episodic memory, executive function, and working memory in a community sample of cognitively asymptomatic older adults. Alzheimer’s & Dementia: Diagnosis, Assessment & Disease Monitoring, 2024. 16(1): p. e12520.

18. Dumurgier, J., et al., Gait speed and decline in gait speed as predictors of incident dementia. The Journals of Gerontology: Series A, 2017. 72(5): p. 655–661.

19. Rudd, K.D., et al., Investigating the associations between upper limb motor function and cognitive impairment: a scoping review. GeroScience, 2023: p. 1–25.

20. Wang, X., et al., Estimating presymptomatic episodic memory impairment using simple hand movement tests: a cross-sectional study of a large sample of older adults. Alzheimer’s & Dementia, 2024. 20(1): p. 173–182.

21. Baudic, S., et al., Executive function deficits in early Alzheimer’s disease and their relations with episodic memory. Archives of clinical neuropsychology, 2006. 21(1): p. 15–21.

22. Rudd, K.D., et al., Stepping and tapping: combining motor tasks improves cognitive classification. GeroScience, 2025: p. 1–14.

23. Rudd, K.D., et al., Hand motor dysfunction is associated with both subjective and objective cognitive impairment across the dementia continuum. Dementia and Geriatric Cognitive Disorders, 2025. 54(1): p. 10–20.

24. Alty, J., et al., Development of a smartphone screening test for preclinical Alzheimer’s disease and validation across the dementia continuum. BMC neurology, 2024. 24(1): p. 127.

25. Rudd, K.D., et al., Investigating the associations between upper limb motor function and cognitive impairment: a scoping review. GeroScience, 2023. 45(6): p. 3449–3473.

26. Gordon, B. and K. Carson, The basis for choice reaction time slowing in Alzheimer’s disease. Brain and Cognition, 1990. 13(2): p. 148–166.

27. Christ, B.U., M.I. Combrinck, and K.G. Thomas, Both reaction time and accuracy measures of intraindividual variability predict cognitive performance in Alzheimer’s disease. Frontiers in human neuroscience, 2018. 12: p. 124.

28. Sano, M., et al., Simple reaction time as a measure of global attention in Alzheimer’s disease. Journal of the International Neuropsychological Society, 1995. 1(1): p. 56–61.

29. Müller, G., et al., Reaction time prolongation in the early stage of presenile onset Alzheimer’s disease. European archives of psychiatry and clinical neuroscience, 1991. 241: p. 46–48.

30. Baddeley, A.D., et al., Attentional control in Alzheimer’s disease. Brain, 2001. 124(8): p. 1492–1508.

31. Gorus, E., et al., Reaction times and performance variability in normal aging, mild cognitive impairment, and Alzheimer’s disease. Journal of geriatric psychiatry and neurology, 2008. 21(3): p. 204–218.

32. Bailon, O., et al., Psychomotor slowing in mild cognitive impairment, Alzheimer’s disease and lewy body dementia: mechanisms and diagnostic value. Dementia and geriatric cognitive disorders, 2010. 29(5): p. 388–396.

33. McGuinness, B., et al., Attention deficits in Alzheimer’s disease and vascular dementia. Journal of Neurology, Neurosurgery & Psychiatry, 2010. 81(2): p. 157–159.

34. Sylvain-Roy, S., L. Bherer, and S. Belleville, Contribution of temporal preparation and processing speed to simple reaction time in persons with Alzheimer’s disease and mild cognitive impairment. Brain and cognition, 2010. 74(3): p. 255–261.

35. Huang, G., et al., Feasibility of computerized motor, cognitive and speech tests in the home: Analysis of TAS Test in 2,300 older adults. The Journal of Prevention of Alzheimer’s Disease, 2025: p. 100081.

36. Alty, J., et al., The TAS Test project: a prospective longitudinal validation of new online motor-cognitive tests to detect preclinical Alzheimer’s disease and estimate 5-year risks of cognitive decline and dementia. BMC neurology, 2022. 22(1): p. 1–13.

37. Bartlett, L., et al., Island Study Linking Aging and Neurodegenerative Disease (ISLAND) Targeting Dementia Risk Reduction: Protocol for a Prospective Web-Based Cohort Study. JMIR research protocols, 2022. 11(3): p. e34688.

38. Kang, M., et al., Subjective cognitive decline plus and longitudinal assessment and risk for cognitive impairment. JAMA psychiatry, 2024. 81(10): p. 993–1002.

39. Cambridge, C., CANTAB Connect Research: Admin Application User Guide v1. 6. Cambridge, UK: Cambridge Cognition Limited, 2019.

40. Lowe, C. and P. Rabbitt, Test\re-test reliability of the CANTAB and ISPOCD neuropsychological batteries: theoretical and practical issues. Neuropsychologia, 1998. 36(9): p. 915–923.

41. Robbins, T.W., et al., Cambridge Neuropsychological Test Automated Battery (CANTAB): a factor analytic study of a large sample of normal elderly volunteers. Dementia and geriatric cognitive disorders, 1994. 5(5): p. 266–281.

42. Brooks, M.E., et al., glmmTMB balances speed and flexibility among packages for zero-inflated generalized linear mixed modeling. 2017.

43. Team, R.C., R: A language and environment for statistical computing. R Foundation for Statistical Computing,. 2021: Vienna, Austria.

44. Australian Bureau of Statistics. Tasmania, 2021 Census All persons QuickStats. 2021.

45. Reisberg, B. and S. Gauthier, Current evidence for subjective cognitive impairment (SCI) as the pre-mild cognitive impairment (MCI) stage of subsequently manifest Alzheimer’s disease. International psychogeriatrics, 2008. 20(1): p. 1–16.

46. Veale, J.F., Edinburgh handedness inventory–short form: a revised version based on confirmatory factor analysis. Laterality: Asymmetries of Body, Brain and Cognition, 2014. 19(2): p. 164–177.

